# Biomimetic 3D mammary duct models of healthy and tumoral tissues engineered by a co-extrusion microfluidic-based technology

**DOI:** 10.64898/2026.04.14.718212

**Authors:** A Richard, V Bergeron, A Boyreau, D Dumousset, E Mazari-Arrighi, G Recher, C Albiges-Rizo, P Nassoy, L Andrique

## Abstract

Engineering the human breast in 3D physio-mimetic models is challenging due to its complex multilayered tubular organization, where milk is produced in acini and transported through ductal structures. These functions rely on a highly organized architecture comprising stromal, epithelial, and extracellular matrix compartments. The dysregulation of this architecture perturbs mammary gland homeostasis and promotes the emergence of diverse breast cancer subtypes, from frequent in situ luminal to rarer metastatic basal-like tumors. Despite this knowledge, conventional anti-cancer drug testing still primarily employs high-throughput 3D spheroid models that account for diffusion but lack stromal components, thereby failing to capture stroma-driven treatment resistance.

With a unique microfluidic co-extrusion platform, we have developed 3D tubular tissues anchored on a porous and biocompatible alginate shell. Using a one-step protocol, we have bioengineered six relevant ductoid models of healthy and tumoral mammary ducts, most notably a multi-layered model comprising a lumen, mammary epithelium, and stromal compartment made of fibroblasts and matrixes. These new models offer limitless applications in tissue engineering including the characterization of an epithelium and its secretory function, and the identification of the stromal influence on healthy and tumoral mammary gland tissue. Finally, by releasing mechanical constraints, we scale-up the tubular duct model into a mammary assembloid that exhibits branching and budding of acini-like structures from the original duct. We envision that this modular design will broadly impact breast basic and clinical research by opening new experimental avenues toward more physio-mimetic tools through the integration of stromal compartments.

## Introduction

The human breast is a complex glandular organ composed of a branching epithelium within a stromal compartment. The functional unit is organized into 15-20 lobes, further divided into lobules and acini and each lobule is connected to the nipple via a duct network^1^. The mammary epithelium itself is characterized by a distinctive bilayer architecture: an inner layer of polarized luminal epithelial cells (EpCs) and an outer layer of contractile basal myoepithelial cells in direct contact with the basement membrane^2,3^. Mammary tissue homeostasis is maintained by an epithelial compartment that is mechanically and biochemically regulated by the surrounding stroma comprising stromal cells like fibroblasts, adipocytes and immune cells embedded within collagen I rich interstitial matrix^4^.

Many 3D models have been developed to replicate solid organs, such as the brain^5^, liver^6^ and spleen^7^. However, just like bronchial epithelium^8^ and artificial vessels^9^, breast epithelium requires the development of complex, functional hollow organs that are more challenging to produce^10,11^. Conventionally, 3D models of the mammary gland are produced using scaffold-based techniques^12^. They often rely on embedding mammary EpCs within Matrigel® that replicates the basement membrane and provide mechanochemical cues to induce cells self-organization in hollow spherical cysts. They rarely exceed 200 µm diameter and 100 µm lumen and moving toward higher scale ductal models (0.5 to 1.2 mm) is essential as it should enhance their biological fidelity and relevance^13–19^. Moreover, the concomitant biofabrication of cystic and ductal topology is still rare because of the need of an anisotropic environment^20,21^. The artificial development of a multi-layered tubular tissue to reconstitute the epithelium, the stromal compartment and the ECM is seldom achieved and requires laborious protocols^22,23^.

In the context of *ex vivo* tumor models, breast cancers (BCs) are typically modelled in 3D by morphologically-limited tumor spheroids^24,25^. Indeed, these simplified models do not allow to discriminate and study all BCs subtypes where interactions between cells and extra-cellular matrix (ECM) are dysregulated and whose origin is from the ducts in 70% of cases^26,27^. BCs comprises a heterogeneous group of pathologies with multiple well-defined subtypes like Luminal and Basal-like that represent 80% of all breast cancers^28,29^ but that have different outcomes and treatments. Luminal BCs are characterized by the epithelial lumen colonization by tumoral cells, and basal-like BCs present invasive tumoral cells that can cross the basal lamina in order to disseminate in other organs. In this context, modeling normal and tumoral mammary gland in 3D need to be optimized and more importantly complexified into an epithelial-stromal tissue with a tailorable technology.

In this study, we developed an original co-extrusion microfluidic-based platform, called the Cellular Capsule Technology (CCT) to biofabricate new 3D ductal mammary models^30^. These models are self-assembled, with a multi-layered complexity, and hosted into tubular, hollow, porous and biocompatible alginate tubes^31^. We successfully developed seven relevant 3D models of the mammary gland and breast cancers that precisely recapitulate the different epithelial-stromal mammary tissue compartments, mammary gland morphogenesis and breast cancers.

## Results

### Biofabrication of a 3D mammary epithelial duct model

To fabricate a mammary hollow epithelium, we employed the CCT (Fig. 1A). Briefly, CCT enables the coextrusion of three distinct solutions through an in-house 3D-printed microfluidic chip: an external sodium alginate solution (ALG), an intermediate solution (IS) of D-sorbitol, and a refrigerated core solution (CS) containing a cell suspension composed of human mammary EpCs (MCF10A) and Matrigel®. The tip of the chip was immersed in a warm calcium chloride bath, inducing rapid gelation of the alginate and Matrigel® solutions into hydrogels^32^. This process yields a hollow tubular structure composed of an external alginate shell and a central mix of EpCs and Matrigel®. Here, cells rapidly adhere to the alginate wall through the matrix mimicking a laminin and collagen IV-rich basement membrane and leads to the formation of a tubular mammary epithelium. We called this first model Mammary Epithelial Ductoid (MED).

**Fig. 1.**
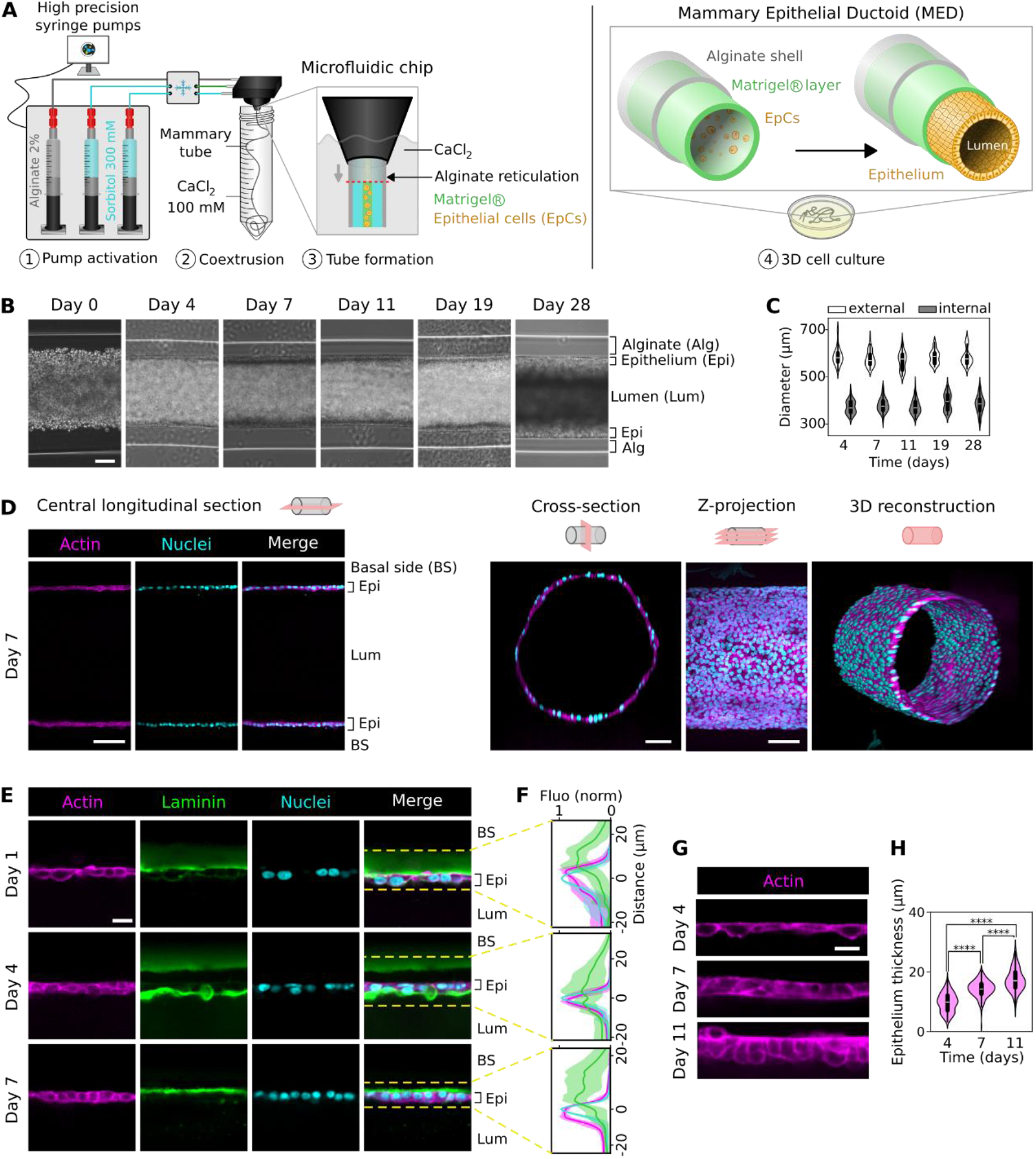
General cellular and matrix architecture of mammary epithelial ductoid (MED) over time. **A**, Schematic representation of CCT experimental setup and steps, and visualization of MED fabrication, formation and maturation. **B**, Bright-field images of MEDs over 28 days of 3D culture. Scale bar, 100 µm. **C**, Quantification of MED’s external (OD) and internal diameters (ID) on days 4, 7, 11, 19, and 28 of culture, from bright-field images (*n* = 16 from 4 distinct images). They correspond to the tissue with (OD) or without (ID) the alginate shell. **D**, Immunofluorescence staining for F-actin (phalloidin, magenta) and nuclei (Hoechst, cyan) on MEDs at day 7. Images represent central longitudinal and cross-sections, a z-stack projection (410 images, z-step = 1.1 µm), and a 3D reconstruction. Scale bars, 100 µm. **E**, Immunofluorescence staining for F-actin (phalloidin, magenta), laminin (green) and nuclei (Hoechst, cyan) on MEDs at day 1, 4 and 28. Images represent the central longitudinal section. Scale bar, 20µm. **F**, Fluorescence intensity profiles for F-actin (magenta), laminin (green) and nuclei (cyan), along the radial axis of the MED’s epithelium at days 1, 4 and 7. For each time point, profiles were generated from confocal images of central longitudinal sections by calculating mean intensities along a line (150 µm wide) from the basal to the luminal side of the epithelium. Fluorescence values were normalized from 0 to 1 based on the minimum and maximum intensities of each line. Distance values were centered at the nuclear peak to measure relative distances between actin, laminin and nuclei. Data represent means values from 10-12 lines across 2-4 different images corresponding to different regions of the MED. **G**, Immunofluorescence staining for F-actin (phalloidin, magenta) on MED’s epithelium at days 4, 7 and 11. Scale bar, 20 µm. **H**, Epithelial thickness measurement at days 4, 7 and 11 from 8 different images (137.5 µm long) along the MED. Within each image, 25 radial measurements were generated between the apical and basal pole of the epithelium. \*\*\*\**P*<0.0001 was determined by Kruskal-Wallis with Dunn’s post-hoc test with Bonferroni correction.

Bright-field images were performed to monitor cellular organization within the tube, from 2 hours (day 0) up to 28 days of culture (Fig. 1B). EpCs were randomly distributed in the lumen and rapidly anchored to the inner wall of alginate. In parallel, the formation of the tubular tissue was observed on a living sample (Supplementary Video 1) and cells exhibited a highly proliferative and migratory phenotype from 2 to approximately 78 hours (∼3 days). Cells proliferated to cover the inner wall of alginate (∼80 hours), and ultimately, proliferation appeared to slow down, likely due to contact inhibition, reaching a confluent state in 3D and a near-complete proliferation arrest by ∼138 hours (5-6 days).

Then we measured the inner and outer diameters (ID/OD) of the MED over time (Fig. 1C). The OD of the MED does not significantly vary, although it differs between separated tubes and along individual tubes, reflecting the intrinsic variability of the CCT. In contrast, the ID presents minor fluctuations, with a mean value increasing from 370.97 µm ± 27.86 µm on day 4, to 383.67 µm ± 36.95 µm on day 28, corresponding to a 3.4% variation. Importantly, at day 6-7, the ID does not significantly vary, which corresponds to cell confluency on the alginate wall, thus to the formation of an epithelium. These measurements are in accordance with the lower end of the physiological range of human lactiferous ducts^19,33^. Taken together, these results show the general stability of the MED for several weeks.

Based on these first observations, we performed actin and nuclei staining on MED at day 7, when the epithelium is formed (Fig. 1D). We were able to image a nice tubular 3D monolayered mammary epithelium on longitudinal and cross-sections of the tubes, and generated a 3D reconstruction (Supplementary Video 2) of the MED model.

To further elucidate cells/ECM organization in greater details, imaging was performed at the beginning of MED formation (Fig. 1E). Actin, nuclei, and laminin fluorescent intensity profiles were measured through epithelial segments (Fig. 1F). On day 1, laminin was localized at the basal side of cells and at intercellular junctions. By day 4, laminin was observed at the luminal side, as well as at the basal side and intercellular junctions. Finally, by day 7, laminin localization was re-established along the basal side of the epithelium. In addition, a partial detachment of the epithelium from the alginate inner wall was observed at day 4, likely due to collective cell contractility. These findings suggest that cells remodel the ECM, coming from Matrigel® source or from their own production, in order to concentrate its deposition on the basal side and maintain cells-ECM interactions. Finally, EpCs morphology was characterized by actin imaging of the MED and epithelium thickness measurement (Fig. 1G-H). On day 4, near-confluent cells exhibited an elongated morphology, while by days 7 and 11, once fully confluent, cells adopted a more cuboidal shape. In parallel, epithelial thickness increased significantly from 9.77 ± 3.19 µm on day 4 to 14.35 ± 2.75 µm on day 7 and 17.35 ± 3.71 µm on day 11.

Together, these results demonstrate the uniqueness of this microfluidic technology to produce a new 3D mammary epithelial model, that is highly reproducible, self-organized and mimicking mammary morphogenesis.

### Mammary Epithelial Ductoid displays epithelial differentiation and low proliferation

MED analysis showed the formation of an epithelium with one (days 4-7) to two (day 11) layers of cells (Fig. 2A and Supplementary Fig. S1A). To evaluate whether this bilayer is organized in two distinct luminal and myoepithelial cells, we performed staining of specific markers as cytokeratin 7 and 17 (CK7/CK17) (Fig. 2A). Results show that both proteins are expressed in EpCs, but no distinct localization in the tissue has been evidenced. This could indicate the absence of luminal/basal orientation in the tissue, corresponding to a partially mature mammary epithelium.

**Fig. 2.**
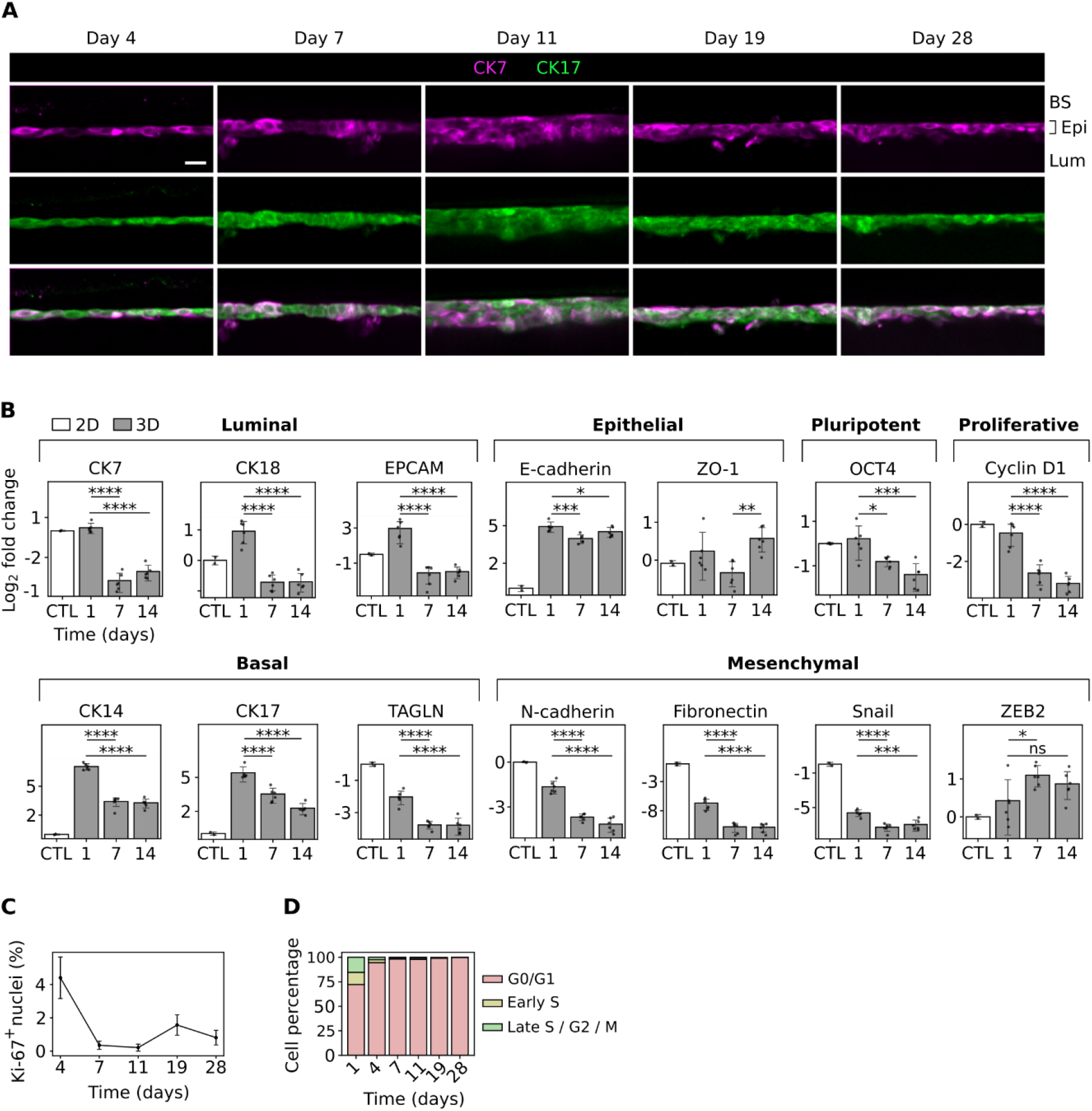
Differentiation, proliferation and cell cycle characterization of the mammary epithelial ductoid (MED) over time. **A**, Immunofluorescence staining for CK7 (magenta) and CK17 (green) on MEDs at days 4, 7, 11, 19 and 28. Images represent the central longitudinal section. Scale bar, 20 µm. **B**, Gene expression analysis for luminal (KRT18/CK18, KRT7/CK7, EPCAM), basal (KRT14/CK14, KRT17/CK17, TAGLN), epithelial (CDH1/E-cadherin, TJP1/ZO-1), mesenchymal (CDH2/N-cadherin, FN1/Fibronectin, SNAI1/Snail, ZEB2), pluripotency (POU5F1/OCT4) and proliferation (CCND1/Cyclin D1) markers on MEDs at days 1, 7 and 14, by RT-qPCR. Transcript levels were normalized to the B2M gene and are presented as fold-change relative to cells in 2D culture (control). Data represent means ± s.d. For 3D conditions, *n* = 6 samples. For 2D control, *n* = 2 samples. \**P*<0.05, \*\**P*<0.001, \*\*\*\**P*<0.0001 were determined by one-way ANOVA with Tukey’s post-hoc test or Kruskal-Wallis with Dunn’s post-hoc test with Bonferroni correction. ns, non-significant. **C**, Proliferative index of epithelial cells on MEDs on days 4, 7, 11, 19, and 28, by automated counting of Ki-67-positive cells and total cells. Counting was performed on *n =* 3 images for each day. **D**, Cell cycle progression assessment by flow cytometry analysis of FUCCI reporter-expressing MCF10A cells in MEDs at days 1, 4, 7, 11, 19 and 28.

Thus, to better characterize proliferation and differentiation state of our tissue we quantified both luminal (KRT7, KRT18, EPCAM) and basal markers (KRT17, KRT14, TAGLN) by RT-qPCR on 2D and 3D culture samples at different time points (Fig.2B). On day 1, both luminal and basal markers are increased in 3D compared to 2D, but ultimately decreased over time. Expression of epithelial (E-Cadherin, ZO1), mesenchymal (N-cadherin, Fibronectin, Snail, ZEB2), pluripotent (OCT-4) and proliferative (Cyclin D1) markers were also analyzed. Interestingly, epithelial markers are overexpressed in 3D cultures of the MED, while mesenchymal markers tend to decrease. In 3D culture, when confluency is reached at day 7, both pluripotency and proliferative markers are downregulated, leading us to hypothesize that an epithelial differentiation occurs.

To confirm the quiescence of our tissue, we measured the ratio of Ki-67 positive and negative cells (Fig. 2C). A significant decline in Ki-67 positive cells was observed between day 4 and 7 of culture, from 4.4 ± 2.77% to 0.35 ± 0.76%, most likely due to contact inhibition as observed in Figure 1. Proliferation remained stable between day 7 and 11, before modestly increasing between day 11 and 19 and stabilizing again until day 28. In parallel, cell cycle progression was measured using modified MCF10A cells expressing the Fluorescent Ubiquitination-based Cell Cycle Indicator (FUCCI) to produce the MED and cell fluorescence was gated by cytometry (Fig. 2D and Supplementary Fig. 1B). The percentage of cells in G0/G1 increased from 72.33% (day 1) to 99.66% (day 28). In parallel, the percentage of early S and late S/G2/M phases respectively decreased from 12.4% (day 1) to 0.32% (day 28), and from 15.3% (day 1) to 0.017% (day 28). A first expansion phase was observed between day 1 and 7, where cells are highly proliferative in order to coat the entire matrix layer and form an epithelium, followed by a non-proliferative epithelial-maturation phase of the tissue. The findings show that MED formation proceeds through two consecutive phases: proliferation and maturation.

### Mammary Epithelial Ductoid displays epithelial polarization and excretory function

The functionality of this MED model was assessed by examining apico-basal polarity and secretory activity, which are intrinsically linked. To assess apicobasal polarity in MED, the Golgi apparatus and nuclei were stained, and fluorescence intensity profiles were analyzed (Fig. 3A-B and Extended Data Fig. 3). On day 4, when EpCs are in a subconfluent state, the Golgi and nuclei intensity peaks overlap. This shows that the Golgi displays a heterogeneous localization around the nucleus, corresponding to a lack of epithelial polarization. From day 7 to day 28, when EpCs are in a confluent state, the Golgi is localized at the luminal side of the cells while the nucleus is localized at their basal side. This demonstrates that EpCs quickly exhibit correct apico-basal polarization, stable for several weeks in culture.

**Fig. 3.**
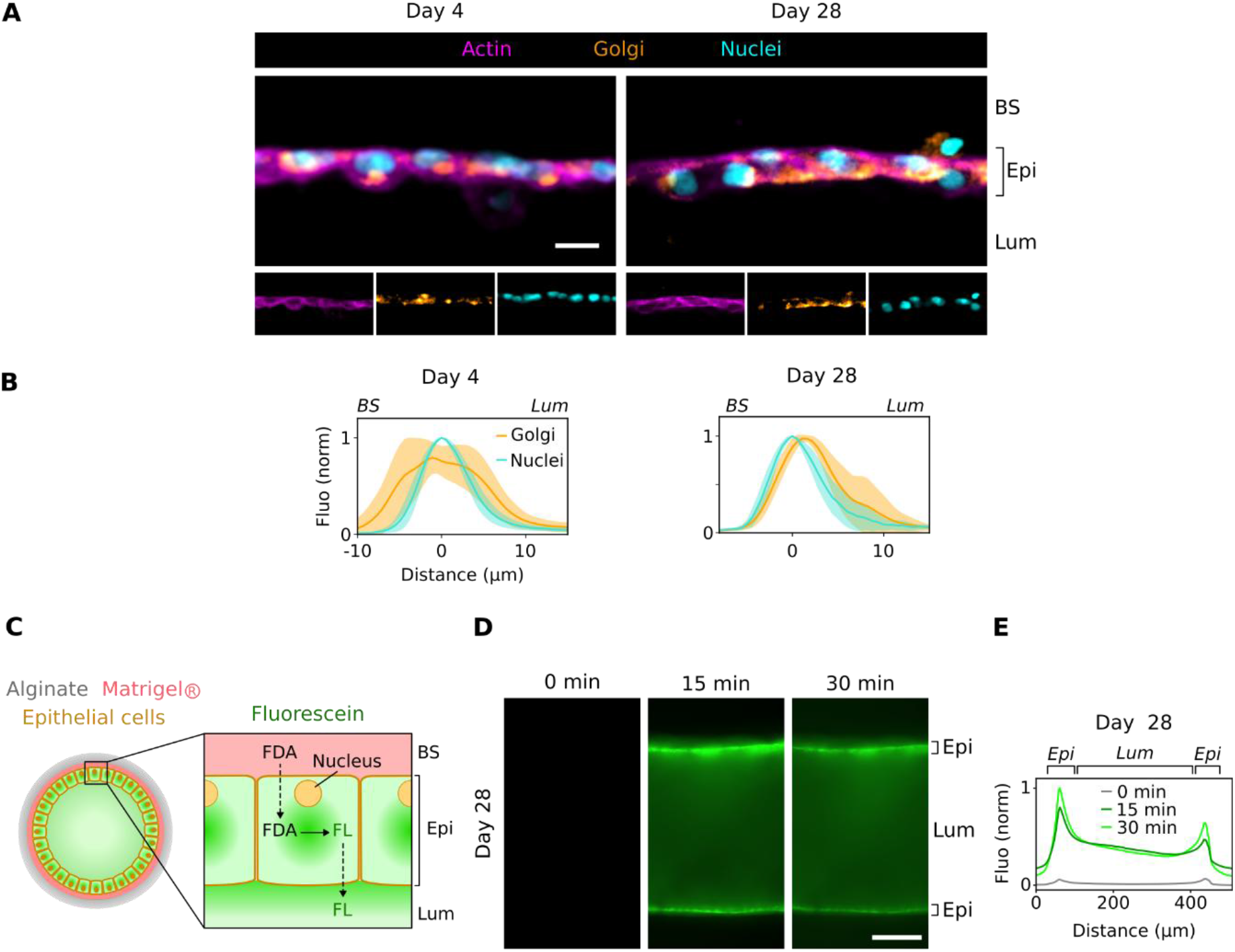
Apico-basal epithelium polarity and oriented excretory function analyses. **A**, Immunofluorescence and staining for F-actin (phalloidin, magenta), Golgi apparatus (Orange) and nuclei (Hoechst, cyan) of MED at days 4 and 28. Images represent the central longitudinal section. Scale bar, 20 µm. **B**, Fluorescence intensity profiles for Golgi (orange) and nuclei (cyan), along the radial axis of the MED’s epithelium at days 4 and 28. For each time point, profiles were generated from confocal images of central longitudinal sections by calculating mean intensities along a line (82.5 µm wide) from the basal to the luminal side of the epithelium (*n* = 10-20, from 2-4 distinct images). Fluorescence values were normalized from 0 to 1 based on the minimum and maximum intensities of each line. Distance values were scaled by assigning the value 0 to the maximum intensity of the nuclei, to measure relative distances between the nuclei and Golgi apparatus. **C**, Schematic representation of the FDA assay in the MED. **D**, Confocal images of fluorescein accumulation in ECs and lumen, taken at the equatorial plane of a 28-days-old MED over 30 minutes. Scale bar, 100 µm. **E**, Fluorescence intensity profiles for fluorescein over a 30 minutes experiment, on day 28. The profile was generated from confocal images at 0 (gray), 15 (dark green) and 30 (bright green) minutes, by calculating mean intensities along a line (165 µm wide) corresponding to a cross-section of the entire width of the MED. Fluorescence values were normalized from 0 to 1 based on the minimum and maximum intensities recorded during the entire assay.

To further evaluate enzymatic activity and excretory capacities of the MED, we performed fluorescein diacetate (FDA) hydrolysis assay (Fig. 3C-E). MED were incubated with non-fluorescent FDA, that should be transformed by esterases into fluorescent fluorescein (FL) and secreted into the lumen of functional EpCs (Fig. 3C). Fluorescence was monitored through live imaging at 0, 15 and 30 minutes after incubation (Fig. 3D) and results are plotted in Fig. 3E. On days 4, 7, 19 and 28, an increase of fluorescence was observed insidethe epithelium after 15 and 30 minutes. This highlights fluorescent accumulation in viable EpCs and functional enzymatic capacities of the MED^34^. In addition, the fluorescence signal observed in the lumen shows that FL is released by apical active efflux^35^. Together, these results indicate that this new biomimetic MED model displays correct apico-basal polarization with an active epithelial luminal excretory function.

### Biofabrication of a multi-layered 3D Mammary Epithelial-Stromal Ductoid (MESD)

To enhance physiological relevance, the MED model was complexified into a bi-layered structure by injecting a mammary stromal-mimetic solution (GFP-fibroblasts, collagen I, Matrigel®) into the microfluidic chip’s intermediate space (Fig. 4A). The resulting coextruded tube is composed of an alginate shell, an intermediate stromal layer (SL) and a central mammary epithelium and was named 3D model Mammary Epithelial-Stromal Ductoid (MESD).

**Fig. 4.**
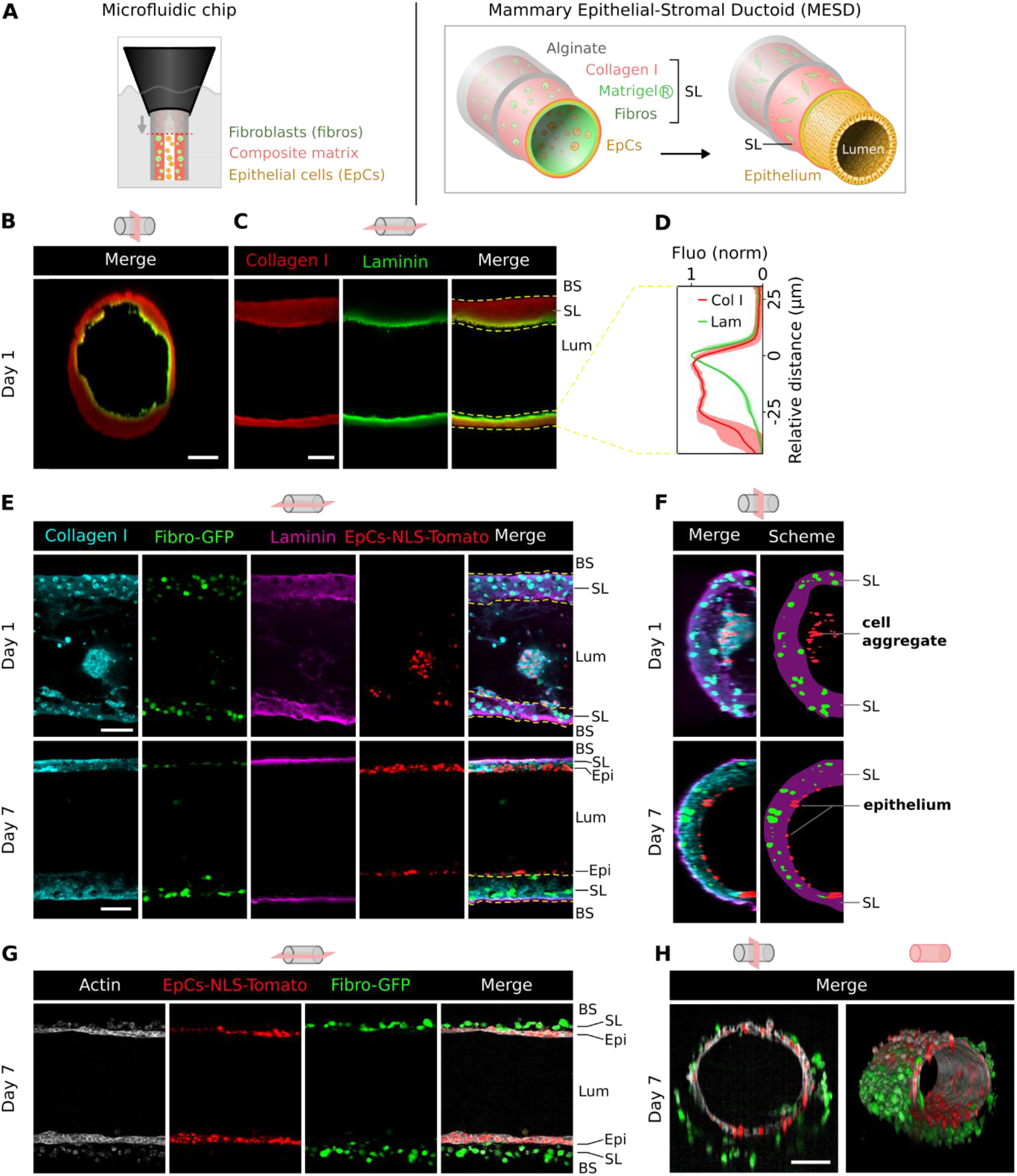
General cellular architecture and matrix analysis of the mammary epithelial stromal ductoid (MESD) over time. **A**, Schematic representation of CCT-based MESD fabrication and formation. **B, C**, Immunofluorescence staining for collagen I (red) and laminin (green) of an acellular variant of the MESD on day 1. Images represent a central longitudinal and cross-section. **D**, Fluorescence intensity profiles for collagen I (red) and laminin (green), along the radial axis of the MESD acellular variant, on day 1. Profiles were generated from confocal images of central longitudinal sections by calculating mean intensities along a line (150 µm wide) from the basal to the luminal side of the tube. Fluorescence values were normalized from 0 to 1 based on the minimum and maximum intensities of each line. Distance values were centered at the laminin peak to measure relative distances between laminin and collagen I. Data represent mean values from 6 lines corresponding to different regions of the tube. **E**-**F**, Immunofluorescence staining for collagen I (cyan), 3T3-GFP fibroblast (green), laminin (magenta) and ECs NLS-Tomato nuclei (red) of MESD on days 1 and 7. Images represent central longitudinal sections and cross-sections. **G, H**, Immunofluorescence staining for F-actin (white), ECs NLS-Tomato nuclei (red) and 3T3-GFP fibroblast (green) on day 1. Images represent a central longitudinal section and a 3D reconstruction. Scale bars, 100 µm.

For a deeper understanding of matrixes gelation and both collagen and laminin spatial organization, we first performed imaging on MESD produced without any cell (Fig. 4B-C). On day 1 a matrix layer with an average thickness of approximately 25 µm was observed. Collagen I seems to be localized at the basal side, while laminin is at the luminal side (Fig. 4D). Together, these results show that the composite matrix organization is similar to the ECM organization of the human lactiferous duct, with basal collagen I mimicking the stromal interstitial matrix, and luminal laminin/collagen IV (Matrigel®) mimicking the basement membrane.

Secondly, MESD was produced with fluorescent cells (GFP-fibroblasts and NLS-Tomato MCF10A cells). Collagen, Laminin, and both GFP/Tomato signaling were observed by confocal imaging, at day 1 and 7 (Fig. 4E-F). On day 1, collagen I and laminin are combined and fibroblasts appear embedded in this intermixed matrix. EpCs are seeded into the lumen with a small amount of matrix to form unorganized aggregates. After 7 days of culture, EpCs proliferated and self-organized into an epithelial tissue anchored to the SL. To better decipher the cellular organization of both EpCs and fibroblasts in MESD, we performed actin staining after 7 days of culture (Fig. 4G-H and Supplementary Video 3). EpCs form a cohesive epithelium while fibroblasts cells are localized at their basal side, embedded in the stromal layer. Together, these results show that we were able to produce a multilayered 3D model self-organized in (1) a basal stromal layer composed of fibroblasts, collagen I, laminin and collagen IV, and (2) a cohesive mammary epithelium laying onto the stroma, recapitulating *in vivo* mammary ducts.

### MESD epithelium displays a luminal and basal orientation

Because fibroblasts are strongly involved in mammary gland morphogenesis^36,37^, we cultured MESD for several weeks in order to observe the epithelium organization in presence of fibroblasts (Fig. 5). Luminal (CK7) or basal (CK17) differentiation markers were localized by confocal microscopy (Fig. 5A) and fluorescence intensity profiles were plotted from the basal to the luminal side of the epithelium (Fig. 5B). On day 4, CK7 and CK17 fluorescence peaks overlap, indicating a lack of proper organization of the two proteins within the epithelium. From day 7 to day 28, CK7 and CK17 fluorescence profiles diverged into two distinct peaks, indicating an organized multilayered epithelium, with CK7 and CK17 being respectively localized at the lumen and basal side. These data imply that fibroblasts rapidly influence tissue-level self-organization of the epithelium with a luminal and a basal layer.

**Fig. 5.**
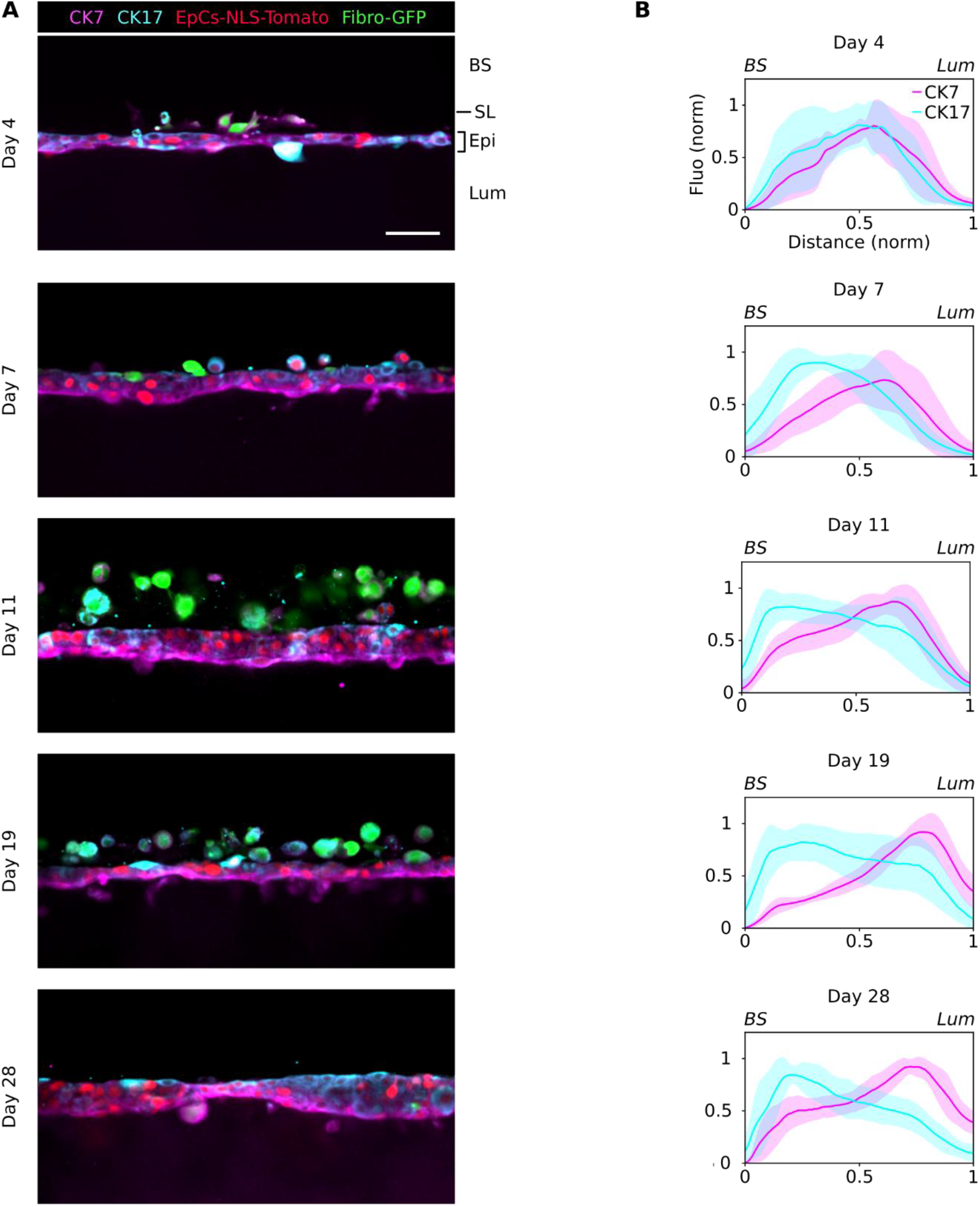
Luminal-basal orientation of the epithelium in the bi-layered mammary epithelial stromal ductoid (MESD). **A**, Immunofluorescence staining for CK7 (magenta), EC nuclei (red), 3T3-GFP fibroblasts (green) and CK17 (cyan) of MESD on days 4, 7, 11, 19 and 28. Images represent the central longitudinal section. Scale bar, 50 µm. **B**, Fluorescence intensity profiles for CK7 (magenta) and CK17 (cyan), along the radial axis of the MESD’s epithelium at days 4, 7, 11, 19 and 28. For each time point, profiles were generated from confocal images of central longitudinal sections by calculating mean intensities along a line (82.5 µm wide) from the basal to the luminal side of the epithelium. To account for variations in epithelium thickness and signal intensity, fluorescence and distance values were normalized and spatially aligned. Profiles were resampled via linear interpolation to a common coordinate system, enabling the calculation of mean intensity distribution for each time point (*n* = 9-14 lines, from distinct images taken along the MESD).

### 3D bioengineering of early luminal and basal-like breast cancers

Subsequently to the development of healthy MED and MESD models, we examined whether the use of tumoral mammary cells instead of healthy mammary cells would result in relevant 3D breast cancer models. To this end, normal EpCs were replaced by luminal MCF7-H2B-GFP or basal-like MDA-MB-231-H2B-GFP cancer cells in both MED and MESD (Extended Data Fig. 6A). In our 3D tubular system, in absence of normal MCF10A cells, luminal MCF7 cancer cells filled the lumen at day 8 when anchored to a laminin matrix, and at day 21 when attached to a stromal layer (Extended Data Fig. 6B). In comparison, basal-like MDA-MB-231 cells, that have more mesenchymal properties, have difficulties to maintain a lumen. More importantly, for both cancer cell types the epithelium is disrupted and not well organized thereby limiting our ability to study tumor cell fate across the epithelium or within the lumen space. Thus, to examine the behavior of these two cancer cell types in a more suited model, tumoral cells were co-cultivated with EpCs with (MESD) and without (MED) the stroma (Fig. 6A-B). Normal MCF10A NLS-Tomato and MCF-7 or MDA-MB-231 H2B-GFP were encapsulated at a ratio of 1-2 cancer cells for 9 normal cells. Here we call these tumoral models Tumoral Mammary Epithelial Ductoid (TMED) and Tumoral Mammary Epithelial Stromal Ductoid (TMESD).

**Fig. 6.**
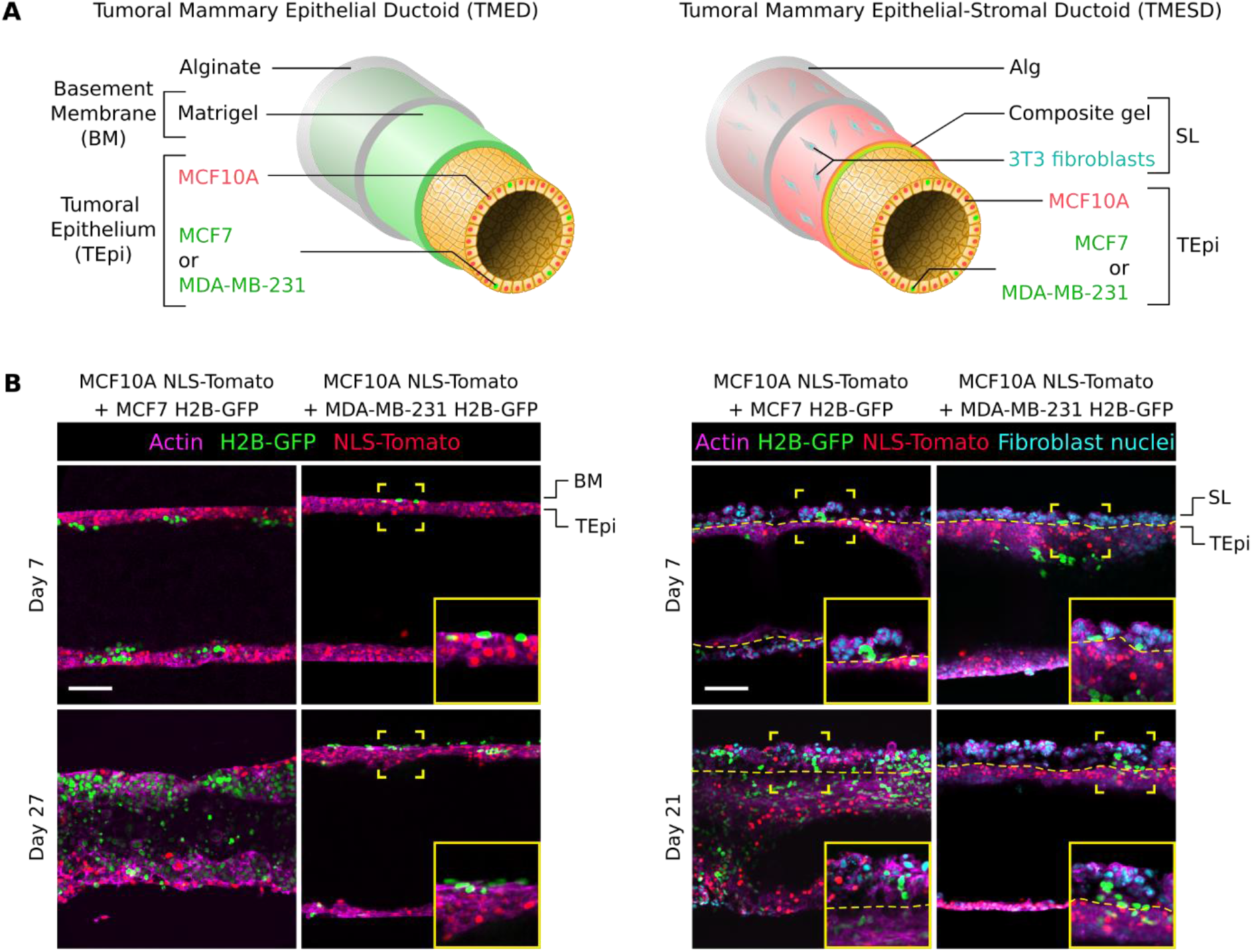
Proliferative and invasive behavior of breast cancer cells in TME(S)Ds in the presence of normal cells. **A**, Schematic representation of TME(S)Ds models in co-culture with normal epithelial cells. **B**, Fluorescence images of actin (phalloidin, magenta), MCF10A-Tomato and MCF7-H2B-GFP or MDA-MB-231 cancer cells (green) when cultivated on MED at days 7 and 27 (left), and MESD at days 7 and 21 (right). Fibroblasts are identified by Hoechst staining. Tubular tissues were produced with 1 or 2 cancer cells for 9 ECs. Yellow brackets indicate the ROI corresponding to the magnified image in the bottom right corner. They highlight different cellular events, like the basal localization of MDA-MB-231 cells in TMEDs, or stromal invasion of MCF7 or MDA-MB-231 cells in TMESDs. Scale bars, 100 µm.

In TMED, luminal MCF7 cells are found at the luminal side of the epithelium at day 7, and filled the lumen after 27 days of culture while basal-like MDA-MB-231 cells rapidly became invasive, transmigrating to the basal side of the epithelium. Interestingly, in TMESD, MCF7 and MDA-MB-231 cells are both capable of invasiveness and although in the luminal side of the epithelium at day 7, they are also found within the stromal layer at day 21. These findings indicate that we were able to develop epithelial and epithelial-stromal models that recapitulate the early establishment and invasiveness of luminal and basal-like breast cancers.

### Bottom-up tissue engineering from a human mammary duct to a mammary gland assembloid

While MESD successfully recapitulates epithelial and stromal compartments of the human mammary gland, it lacks terminal end buds (TEBs), which are bulb-shaped structures also called acini, that arise from the duct and invade the surrounding stroma. These TEBs are active growing regions of the mammary gland that allow the monitoring of mammary morphogenesis. In our MESD model, the stromal layer is approximately 25 µm thick, which might prevent cell proliferation, matrix invasion, branching and budding.

To overcome this issue, we scale-up MESD: after 7 days of culture, the alginate shell was removed, and epithelial-stromal ductoids were included in 300 µl of a composite matrix composed of type I collagen and Matrigel® (Fig. 7A). Branching of the ductoid in numerous bulb-shaped structures was revealed by nuclei and actin staining after seven days of culture of the assembloid (Fig. 7B.a-b).

**Fig. 7.**
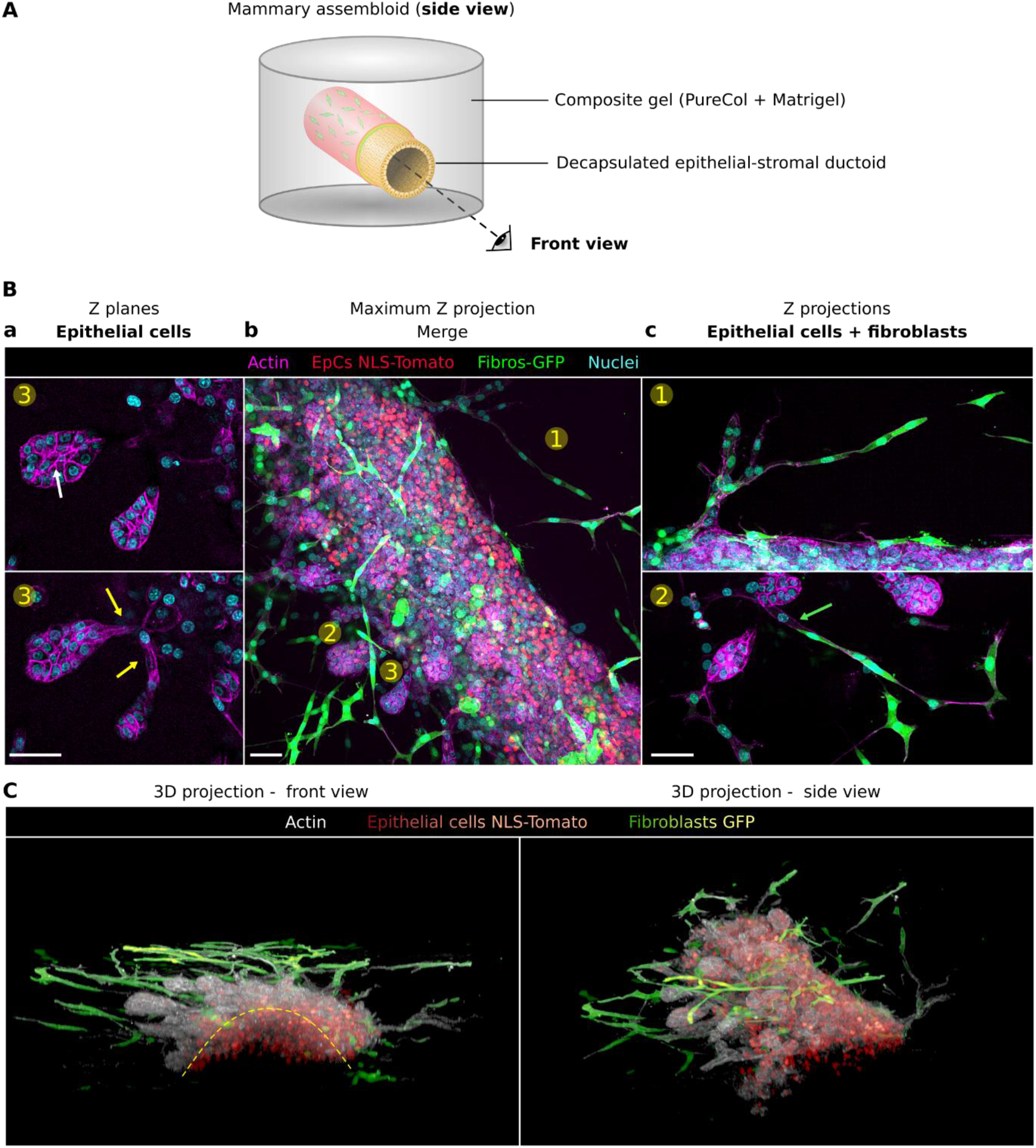
Branching and budding capacities of the mammary assembloid model. **A**, Schematic representation of the experimental design of the mammary assembloid. **B**, Immunofluorescence and staining for F-actin (phalloidin, magenta), ECs NLS-Tomato nuclei (red), GFP fibroblasts (green) and nuclei (Hoechst, cyan) of the mammary assembloid at day 6. (**a**) Branching and budding ECs emerging from the main ductal structure (**b**). (**c**) Highlights on fibroblasts spreading. The white arrow shows the formation of a micro-lumen at the center of a bud, yellow arrows indicate branching structures connecting the main duct to buds, and the green arrow pinpoints the physical interactions between a fibroblast and a bud. Scale bars, 50 µm. **C**, Immunofluorescence and staining for F-actin (phalloidin, white), ECs NLS-Tomato nuclei (red) and 3T3-GFP fibroblasts (white) of the mammary assembloid at day 6. Images represent front and side views of a full z-stack 3D projection (300 images, z-step = 0.308 µm).

These structures are connected to the epithelium of the original duct and mimic the ductal elongation and branchingofamammary gland, suggesting TEBs in formation (Fig. 7B.a). Furthermore, branching and duct elongations were correlated with stretched fibroblasts accumulation surrounding the tip of the new bulb-shaped structures (Fig. 7B.c). Of note, the development of such complexity does not disrupt the overall architecture of the mammary epithelium as a hollow organ (Fig. 7C). Finally, releasing mechanical constraints enabled the tubular duct model to develop into a mammary assembloid displaying branching and budding of acini-like structures.

## Discussion

Replicating the epithelial–stromal architecture of the mammary gland remains a major challenge for improving the biological relevance of *in vitro* breast models. Such biomimetic systems would enable investigation of the individual and combined roles of epithelial cells, stromal components, and the extracellular matrix during the earliest stages of cancer initiation. Capturing these spatiotemporal tissue dynamics is critical for uncovering new mechanisms of mammary homeostasis and tumorigenesis and for advancing diagnostic and therapeutic strategies.

Our unique CCT offers a streamlined approach to tissue assembly and allows us to generate versatile 3D tubular mammary ductoid models which can be implemented on demand. In this study, we engineered six new 3D models of healthy and tumoral mammary ductoids and one mammary gland assembloid recapitulating the complex morphogenesis of the mammary gland (Fig. 8).

**Fig. 8.**
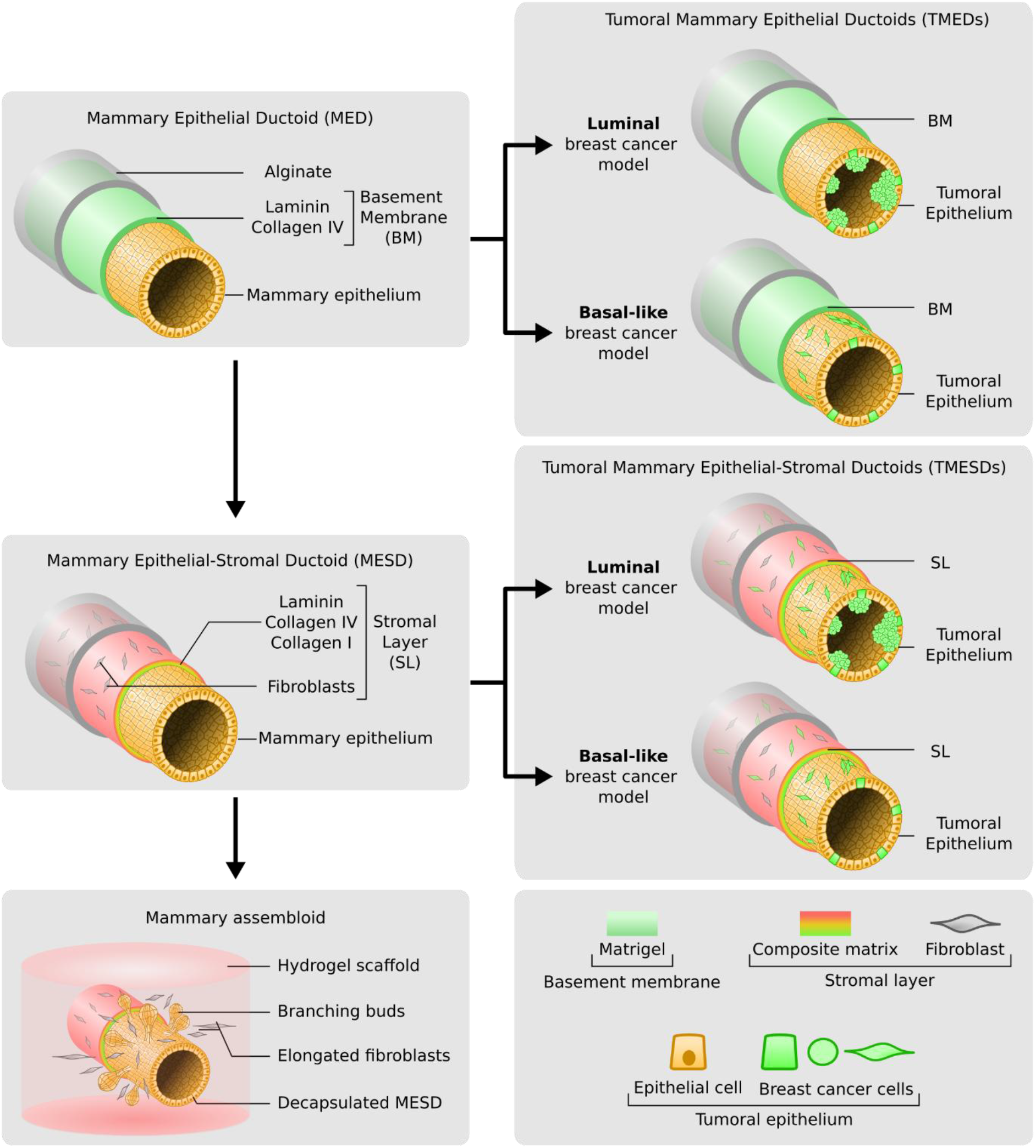
Modular 3D platform for mammary gland and breast cancer models. Schematic representation of the experimental design for the biofabrication of 3D in vitro models of the mammary gland and breast cancer.

The first MED model was developed to mimic a mammary epithelium anchored to a basement membrane, and is the benchmark model to elucidate epithelial cell-cell interactions, apico-basal orientation, and excretory capacity in the lumen. To go further, we were able to implement the MED by incorporating the stromal layer: fibroblasts, collagen I and laminin. For the first time and in one step, we imposed an epithelial-stromal architecture that quickly self-organized in a multilayered and organized tissue. Here we engineered human mammary ductoids with physiological diameter, and meter-scale length that offers rare events observation and enhances statistical robustness.

By adding a stromal layer in the second MESD model, we successfully managed to model a functional microenvironment. Under the influence of both fibroblasts and ECM on the tissue, EpCs undergo luminal and myoepithelial differentiation and cancer cells invade the stroma. This is in accordance with stromal invasive behavior of luminal MCF7 cancer cells observed in the literature^38^ and demonstrating the relevance of the fibroblasts in TMESD model.

Here we demonstrated that ductoid models, when encapsulated in an alginate tube and with a collagen I rich stroma, display low proliferative phenotype similar to mature mammary ducts and we hypothesized that mechanical constraint of the alginate shell inhibits cell proliferation^12,39,40^. So, in order to mimic the mammary gland tissue with both non-proliferative ducts and proliferative acini (TEBs), we removed the alginate shell before inclusion in a collagen/laminin higher-scale scaffold. As expected, EpCs entered in a proliferative state to form branching/budding acini-like structures from the duct, and fibroblasts have spread into the matrix in a more elongated morphology. We propose that the laminin and collagen IV ECM of the assembloid model drives EpCs lobular morphogenesis to form proliferative TEBs. Finally, the presence of elongated fibroblasts along these TEBs structures show the importance of a complex stroma with fibroblasts that guide ductal elongation, branching geometry, and epithelial invasion through biochemical and mechanical signaling^36,37,41^.

In BCs, cancer associated fibroblasts (CAFs) are also key regulators of tumor progression and were identified in therapy resistance^38,41^. However, robust evaluation of tumorigenic potential still relies on *in vivo* experimentation because of the difficulty to reproduce stromal–epithelial interactions *in vitro*, particularly those mediated by fibroblasts. We postulate that a tumoral assembloid model would be relevant to decipher fibroblasts’ pivotal role on both healthy and tumoral processes of the mammary gland. Finally, as adipocytes, endothelial and immune cells have been described as major regulators of physiological and pathological processes of the mammary gland, all our 3D models offer limitless possibilities for customization. In conclusion, with our seven sophisticated 3D models we developed a unique, robust and evolutive platform for mammary gland tissue engineering associated with morphogenesis and invasion studies and broadly applicable to diverse tissue models.

## Methods

### Cell culture and genetic modifications

In 2D culture, MCF10A cells were cultured in Dulbecco’s Modified Eagle Medium (DMEM)/F-12 without phenol red (Gibco) supplemented with 10% Foetal Bovine Serum (FBS; Capricorn Scientific), 1% penicillin-streptomycin (P-S; Gibco), 10 µg/ml insulin (Gibco), 20 ng/ml human Epidermal Growth Hormone (hEGF; Gibco), 100 ng/ml cholera toxin (Sigma) and 0.5. µg/ml hydrocortisone (Sigma). NIH-3T3, MCF7 and MDA-MB-231 cells were cultured in high glucose DMEM (Gibco) supplemented with 10% FBS, 1% P-S. All cells were passaged using 0.05% trypsin (Gibco). All cells were maintained in culture at 37°C, 5% CO2, and fed every 3 days. In 3D culture, all mammary ductoid models were maintained in the same media as MCF10A cells, with lower concentration of hEGF at 1 ng/ml and the addition of 10 µM Y-27632 ROCK inhibitor (Stemcell). Half of the conditioned media volume was replaced with fresh media every 3 days.

To visualize their nuclei, MCF10A cells were transduced with a custom-engineered lentiviral vector expressing TdTomato-NLS (Nuclear Localization Sequence) fusion protein under the constitutive control of a CMV (Cytomegalovirus) promoter. Similarly, nuclear-labeled MCF-7 and MDA-MB-231 cell lines were generated using a custom construct using eGFP fused to histone H2B, under the control of a UbiC (human ubiquitin C) promoter. This vector integrated a puromycin-resistance cassette to facilitate the selection of transduced cells.

For cell cycle monitoring, MCF10A cells were transduced with the commercial pBOB-EF1-FastFUCCI-Puro plasmid^42^ (Addgene #86849). It comprises a dual-color reporter consisting of mKusabira-Orange 2 (mKO2) fused to hCdt1 (G1 phase) and mAzami-Green 1 (mAG1) fused to hGeminin (S/G2/M phases), under the control of an EF1 promoter a puromycin-resistance cassette to facilitate transduced-cells selection.

### Microfluidic fabrication of 3D mammary ductoids

The Cellular Capsule Technology (CCT), as previously described^31^, allows the encapsulation of cells and matrix in a hollow alginate tube. In the original version of CCT, three distinct solutions are coextruded inside an in-house 3D printed microfluidic chip: an external solution (ALG) of 2% (w/v) sodium-alginate (AGI, I3G80), an intermediate solution (IS) of 300 mM D-sorbitol (Sigma), and a core solution (CS) composed of cells with or without matrix. The flow rates used were: 5 ml/hour for the ALG solution and 2.5 ml/hour for IS and CS solutions. The tip of the chip was immersed in a 50 ml tube containing a 100 mM CaCl2 solution (Sigma), previously heated at 37°C, to allow rapid gelation of alginate and matrix hydrogels. While all models used 2% alginate as their ALG solution, IS and CS solutions composition varied as detailed below.

MED were produced using 300 mM D-sorbitol as IS, and a mix of MCF10A cells, Matrigel® and DMEM/F-12 without phenol red at a final volume ratio of 1:2:2. MESD were produced following the same protocol as for MED, but D-sorbitol was replaced in IS by a solution of 10 µl of 3T3-GFP fibroblasts, 15 µl of 0.67% (w/v) alginate, 25 µl of PureCol® EZ Gel (1.25 mg/ml final) and 50 µl of Matrigel® (50% v/v). A cell-depleted variant of the MESD was fabricated by replacing cell-volume by DMEM/F12 without phenol red. For TMED and TMESD production, MCF10A were replaced by a mix of MCF10A-NLS-Tomato and MCF-7 H2B-GFP or MDA-MB-231 H2B-GFP in the CS, at a ratio of 9 normal cells for 1-2 tumoral cells. All 3D models were developed between the VoxCell core facility (TBMCore, UAR CNRS 3427, Inserm US005, University of Bordeaux) and the LP2N/BiOf team.

### Mammary assembloid fabrication

In order to fabricate the mammary assembloid model, seven days old MESD was manually cut into approximately 1-2 cm long segments. Then, the alginate shell was dissolved by incubating the segments in a 0.1 mg/ml alginate lyase (Sigma) solution at room temperature for a few minutes, under real-time microscopy observation. In parallel, 300 µl of composite gel, consisting of 1 volume of PureCol® EZ Gel (1.25 mg/ml), 2 volumes of Matrigel® and 1 volume of DMEM, was poured in a Petri dish and gelled at 37°C for a minimum of 30 minutes. Next, tissue segments were harvested and placed onto the composite gel. Subsequently, the same volume of composite gel was poured onto the segments before gelation at 37°C during 30 minutes. Finally, embedded tissue segments were incubated in media and cultured in the same way as the mammary ductoids for six days.

### RT-qPCR

Total RNA was extracted from 2D culture of MCF10A cells at 70% of confluence, and from MEDs segments at different culture stages (days 1, 7 and 14). For 3D MED sections, alginate shell and Matrigel® were dissolved using ReLeSR™ reagent at room temperature for a few minutes under microscopy observation, before being rinsed with PBS 1X (Corning) before cell lysis. Total RNA was purified using the Nucleospin RNA mini kit (Macherey-Nagel) and treated with RNAse-free DNAse (Macherey-Nagel). Subsequently, 500 ng of total RNA was reverse transcribed into cDNA using Superscript™ IV First-Strand Synthesis System (Thermo Fisher Scientific).

RT-qPCR were performed at the OneCell facility (TBMCore, UAR CNRS 3427/Inserm US 005), using GoTaq(R) qPCR Master Mix (Promega) and gene-specific primers (Supplementary Table 1) on the CFX 384 (Biorad). The cycling parameters were as follows: initial denaturation (95 °C, 15 s), followed by 45 cycles of amplification and SYBR signal detection (95 °C denaturation, 15 s; 60 °C annealing, 30 s; 72 °C extension, 30 s). For data analysis, gene expression was normalized to B2M expression as the reference gene, with 2D culture sample serving as the control (n=2). For each gene of interest and time point, six biological replicates (n=6) were analyzed. Relative expression levels were determined using the 2^ddCt method. For data visualization, results were presented as fold-change relative to the control (2^ddCt). However, to ensure statistical validity, dCt values were used for statistical analyses because they better reflect the normal distribution of the replicated data.

### Confocal immunofluorescence imaging

Tissues segments were fixed with a 4% formaldehyde solution at RT for 2h. Saturation, permeabilization and primary antibody incubation were performed simultaneously with a solution of DMEM (Pan Biotech) with 2% FBS, 1% BSA (Pan Biotech), 0.1% Triton-X100 (Applichem), and 1/500 antibody solution, at 4°C for 24h. After being rinsed with DMEM, tissue segments were incubated with secondary antibodies, phalloidin and hoechst dyes, all diluted at 1/1000 in DMEM, at 4°C for 12h. The primary and secondary antibodies used are listed in Supplementary Table 2.

Samples were optically cleared with 200 µl of the FUnGI clearing agent^43^ at RT overnight. Once cleared, tissue segments were mounted with FUnGI in 0.5 mm thick spacers to be imaged using a spinning disk confocal microscope (Leica DMi8 Yokogawa CSU). The microscopy was done at the Bordeaux Imaging Center, a service unit of the CNRS-INSERM and Bordeaux University. Mammary assembloids were fixed and cleared in the same way, and imaged using an Andor BC43 confocal microscope (Oxford Instruments Andor, Belfast, UK) equipped with 405, 488, 561 and 638 nm lasers for illumination. Z-stack images were acquired with a 20 × 0.8 NA objective lens collected at 0.65 µm step size.

### Proliferation index measurements

To quantify the proliferative index of the MED over time, confocal imaging was conducted on tissue segments fixed at days 4, 7, 11, 19 and 28 of culture. Samples were immunostained for Ki-67 and counterstained with Hoechst to visualize total nuclei. Then, using a home-made ImageJ automated command, automated Ki-67-positive nuclei and total nuclei counting was performed (n = 3 images for each day), to assess the percentage of proliferating cells at each time point.

### Flow cytometry and cell cycle analysis

3D samples (MED segments) were decapsulated prior to cell dissociation, using ReLeSR™ reagent at room temperature for a few minutes under microscopy observation, before washed in DMEM. 2D and 3D samples were dissociated using 0.05% trypsin (Gibco) before being fixed using a 4% PFA solution at room temperature for 5 minutes and resuspended in PBS. Non-stained MCF10A cells from 2D culture were used as a control to subtract nonspecific background noise. Flow cytometry experiments were performed at the FACSility facility (TBMCore, UAR CNRS 3427/Inserm US 005) using the BD LSRFortessa™ Cell Analyzer. Data was analysed using FlowJo™ v10. Gating strategies are indicated in the supplementary figure 1-B.

### FDA functional assay

Fluorescein diacetate (FDA) was diluted, at a final concentration of 6.25 µl/ml, in a solution of DMEM/F-12 without phenol red supplemented with 1% P-S, 10 µg/ml insulin, 1 ng/ml hEGF, 100 ng/ml cholera toxin and 0.5. µg/ml hydrocortisone. MED segments were incubated in this FDA solution for 5 minutes at 37°C with 5% CO2, before being rinsed 3 times with the same solution depleted of FDA. Then tissue segments were imaged with a confocal spinning disk microscope (Nikon Eclipse Ti2-E) at the central longitudinal plane for 30 minutes. Images taken at 0, 15 and 30 minutes were analysed on ImageJ, where fluorescence intensity profiles were measured along a line (165 µm wide) corresponding to a cross section of the entire width of the MED. Fluorescence values were normalized from 0 to 1 based on the minimum and maximum intensities recorded during the entire assay.

### Histology

MED samples were fixed with 4% PFA at room temperature for 2h. Then, haematoxylin-oesin (HE) staining was performed using standard protocols. Subsequently, tissue segments were first embedded in 2% agarose (Thermo Fisher Scientific), then in paraffin and finally cut into 3 µm thick sections, followed by observation under bright-field microscopy (Leica RM2255). Histology experiments were performed at the Histopathologie facility (TBMCore, UAR CNRS 3427/Inserm US 005).

### Image analysis

Image processing and quantitative analysis were performed using ImageJ/Fiji software, with the exception of the 3D projection of assembloids which were generated using Imaris software.

To track changes in MED dimensions over time, internal (ID) and external diameters (OD) were measured using bright field microscopy images of MEDs at six different culture time points (days 0, 4, 8, 11, 17 and 28). OD/ID respectively corresponds to the diameter including the alginate shell, or not. Three independent MEDs were analyzed, and four representative regions of interest (ROIs) were selected for each time point. Four measurements were obtained manually for each ROI.

To determine cellular and matricial organization within MEDs over time, confocal microscopy images were taken at the equatorial plane of MED segments, stained for actin, laminin and nuclei, at day 1, 4 and 7. For each time point, fluorescence intensity profiles were measured from two or four images by tracing 10-12 lines (150 µm wide) along the radial axis of the epithelium. Fluorescence values were normalized from 0 to 1 based on the minimum and maximum intensities of each ROI. Distance values were scaled by assigning the value 0 to the maximum intensity of the nuclei, to measure relative distances between the actin, nucleus and laminin.

To assess epithelium thickness changes over time, confocal microscopy images were taken at the equatorial plane of actin-stained MED segments at day 4, 7 and 11. For each time point, eight images (137.5 µm in length) were analyzed. Within each ROI, 25 radial measurements were taken between the basal and luminal poles of the epithelium. Abnormal values were manually identified and removed.

To evaluate the apico-basal polarization of the MED over time, confocal imaging was performed after immunostaining of the Golgi apparatus combined with nuclear staining of fixed tissues at different time points (days 4, 7, 11, 19 and 28). For each time point, 1-4 images were taken at the equatorial plane of the MED, on which 7-20 lines (82.5 µm wide) were traced from the basal to the luminal side of the cells. Fluorescence intensity profiles of the Golgi apparatus and nuclei were measured along each line, and fluorescence values were normalized from 0 to 1 based on the minimum and maximum intensities of each line. Distance values were scaled by assigning the value 0 to the maximum intensity of the nuclei, to measure relative distances between the nuclei and Golgi apparatus.

To investigate the luminal-basal localization of laminin and collagen I within the MESD, a cell-depleted variant of it was fabricated (as previously described) and labeled for both proteins. Then confocal imaging of a segment fixed on day 1 was conducted. An image of the cell-free MESD was taken at the equatorial plane, on which six different lines were traced along the radial axis. Fluorescence intensity profiles of the laminin and collagen I were extracted from each line and values were normalized from 0 to 1 based on the minimum and maximum intensities of each line. Distance values were scaled by assigning the value 0 to the maximum intensity of the laminin signal, to measure relative distances between the laminin and collagen I.

To observe the luminal-basal distribution of CK7 and CK17 within the epithelium of the MESD over time, confocal imaging was performed after immunostaining of both proteins on tissues fixed at different time points (days 4, 7, 11, 19 and 28). For each time point, 1-2 images were taken at the equatorial plane of the MESD, on which 9-14 lines (82.5 µm wide) were traced from the basal to the luminal side of the cells. Fluorescence intensity profiles of CK7 and CK17 were measured along each line. Both fluorescence and distance values were normalized to account for variability in fluorescence signal and epithelium thickness. Because the physical length of the ROIs varied, the normalized distance coordinates were resampled via linear interpolation to a uniform interval. This standardization facilitated the alignment of all profiles onto a common coordinate system, enabling the generation of a representative mean intensity plot for each condition.

### Statistics

Statistical analyses were performed in Python (v.3.8.19) using SciPy (v.1.10.1), Statsmodels (v.0.14.1) and Scikit-posthocs (v.0.8.1) libraries. Data were structured using Pandas. For all data sets, normal distribution and homogeneity of variances were checked using respectively Shapiro-Wilk and Levene’s tests. In the case of normally distributed data and equal variances, one-way ANOVA test was performed combined with Tukey’s HSD post-hoc test to compare each condition with each other. If the data fails the normality or variance tests, Kruskal-Wallis test was performed combined with Dunn’s post-hoc test with Bonferroni correction. A p-value inferior to 0.05 was considered statistically significant.

## Acknowledgements

We thank all members of the BioImaging and OptoFluidics team for joining discussions on this project. We extend our gratitude to the LP2N UMR CNRS 5298 and the IOGS who gave us the opportunity to work in a multidisciplinary environment. We acknowledge the TBMCore unit, a CNRS-Inserm and Bordeaux University unit, and all the facilities for their contribution to the different experiments: VoxCell, FACSility, OneCell and Histopathologie. We thank Atika Zouine for her help on cytometry analysis. We thank the Bordeaux Imaging Center, a service unit of the CNRS-INSERM and Bordeaux University, member of the national infrastructure France BioImaging supported by the French National Research Agency (ANR-10-INBS-04), and we acknowledge the involvement of Magali Mondin, Sébastien Marais and Jérémie Teillon. This project was supported by La Ligue Contre le Cancer Gironde and the INCA PL-BIO. We are grateful to the national Inserm booster program Mecacell 3D, the Inserm ITMO cancer PCSI and the ANR MATISSE for V.B. and A.R. salaries.

## Author contributions

A.R., L.A., C.A.R., G.R., P.N. contributed to the conception of the study. A.R. and L.A. designed the research. A.R., V.B., A.D.B., D.D., E.M.A., L.A. carried out all experiments and performed image analysis. A.R. generated figures and statistical analysis. A.R. and L.A. drafted the paper. All authors provided feedback on the paper.

---

**Extended Data Figure 3.**
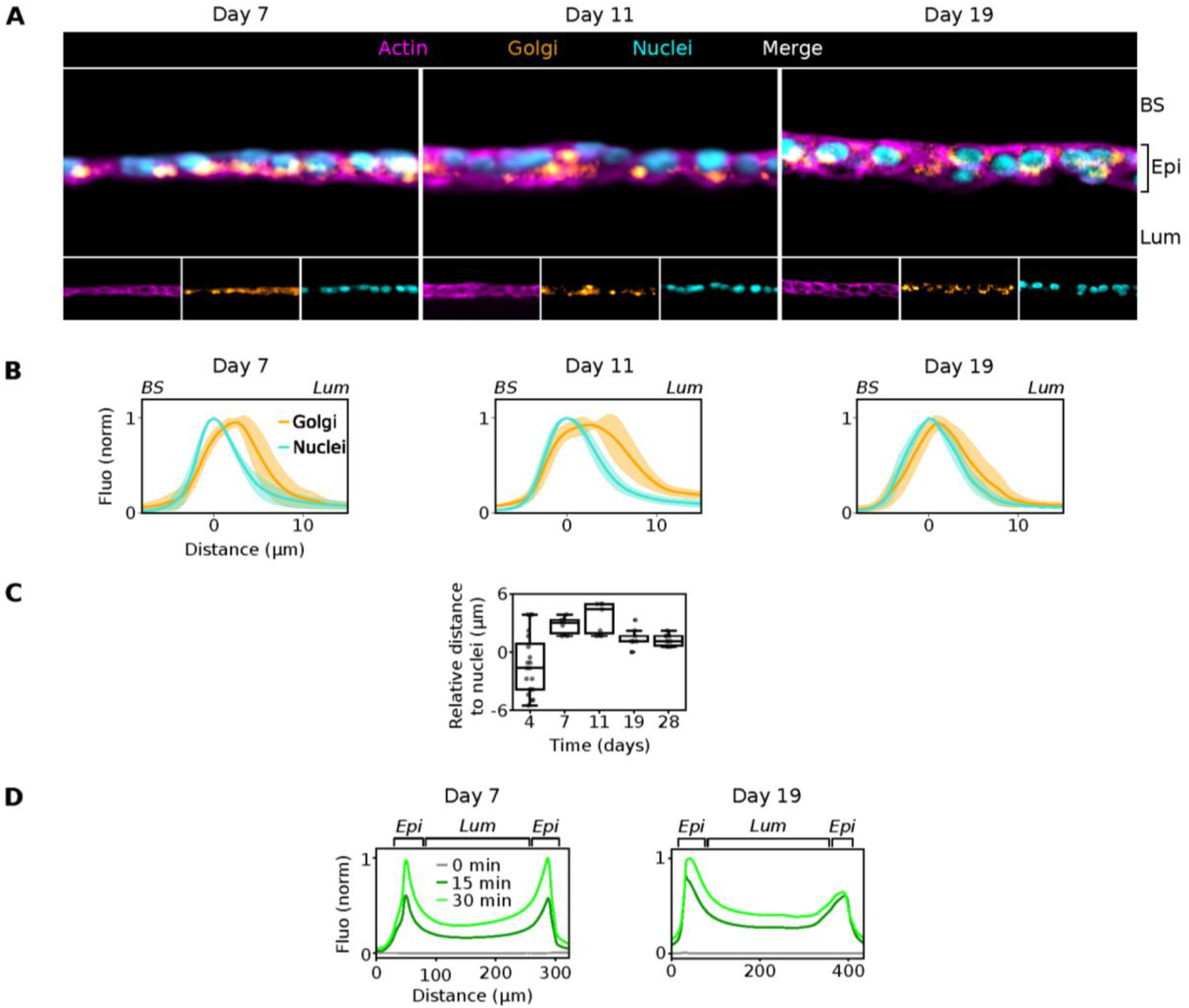
Apico-basal epithelium polarity and oriented excretory function analyses. **A**, Immunofluorescence and staining for F-actin (phalloidin, magenta), Golgi apparatus (Orange) and nuclei (Hoechst, cyan) of MED at days 7, 11 and 19. Images represent the central longitudinal section. Scale bar, 20 µm. **B**, Fluorescence intensity profiles for Golgi (orange) and nuclei (cyan), along the radial axis of the MED’s epithelium at days 7, 11 and 19. For each time point, profiles were generated from confocal images of central longitudinal sections by calculating mean intensities along a line (82.5 µm wide) from the basal to the luminal side of the epithelium (*n* = 6-9, from 1-3 distinct images). Fluorescence values were normalized from 0 to 1 based on the minimum and maximum intensities of each line. Distance values were scaled by assigning the value 0 to the maximum intensity of the nuclei, to measure relative distances between the nuclei and Golgi apparatus. **C**, Relative distance between Golgi apparatus and nuclei in MEDs at days 4, 7, 11, 19 and 28. **D**, FDA experiment’s fluorescence intensity profiles for fluorescein over a 30 minutes experiment, on day 7 and 19. Profiles were generated from confocal images at 0 (gray), 15 (dark green) and 30 (bright green) minutes, by calculating mean intensities along a line (165 µm wide) corresponding to a cross-section of the entire width of the MED. Fluorescence values were normalized from 0 to 1 based on the minimum and maximum intensities recorded during the entire assay.

**Extended Data Figure 6.**
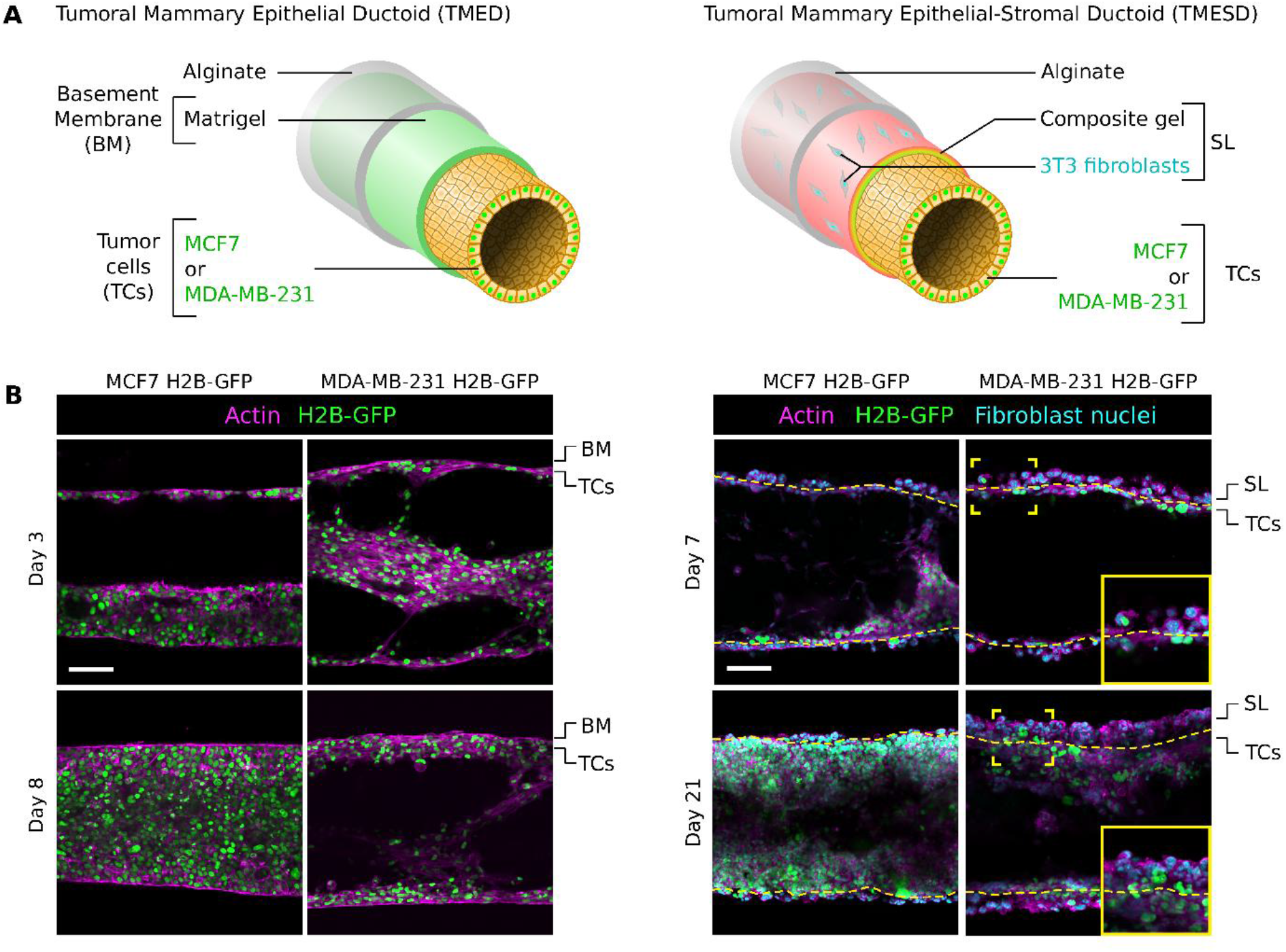
Proliferative and invasive behavior of breast cancer cells in TME(S)Ds in the absence of normal cells. **A**, Schematic representation of TME(S)Ds models in absence of normal epithelial cells. **B**, Fluorescence images of actin (phalloidin, magenta) and MCF7-H2B-GFP or MDA-MB-231 cancer cells (green) when cultivated on MED at days 3 and 8 (left), and MESD at days 7 and 21 (right). Fibroblasts are identified by Hoechst staining. Yellow brackets indicate the ROI corresponding to the magnified image in the bottom right corner. They highlight different cellular events, like the stromal invasion of MCF7 or MDA-MB-231 cells in TMESDs. Scale bars, 100 µm.

