## supplementary figure and tables for "Biomimetic 3D mammary duct models of healthy and tumoral tissues engineered by a co-extrusion microfluidic-based technology"

**Supplementary Figure S1**

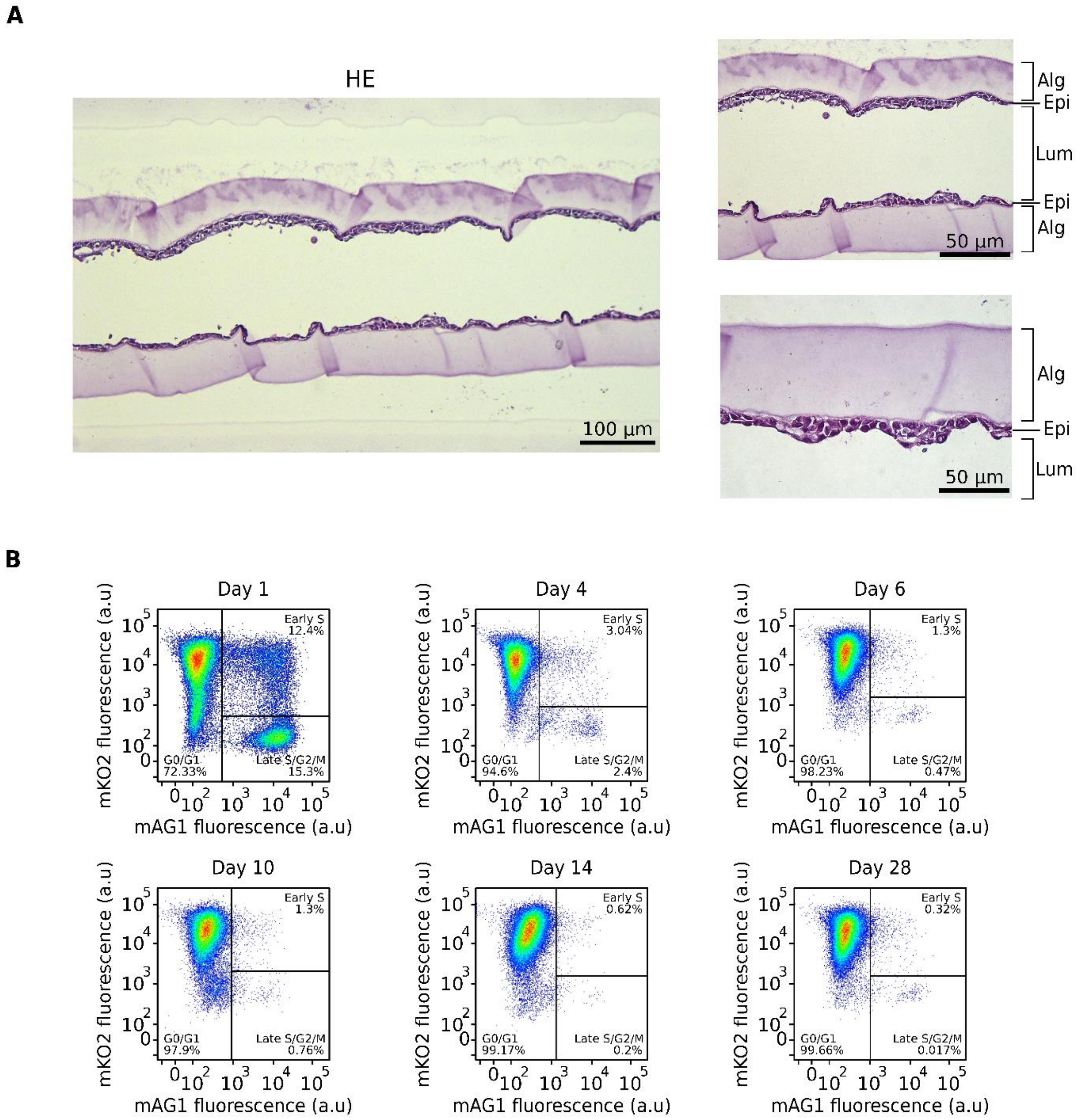


**Supplementary Fig.S1 | Histology and cell cycle analyses of the MED.**
**A**, Bright-field images of HE-stained MED at day 11. **B**, Cell cycle progression assessment by flow cytometry analysis of FUCCI reporter-expressing MCF10A cells in MEDs on days 1, 4, 7, 11, 19 and 28.

**Supplementary Table 1**

| Primary antibodies | | | | |
|---|---|---|---|---|
| **Target** | **Host** | **Company** | **Cat. N°** | **Dilution** |
| Laminin | Rabbit | Abcam | ab11575 | 1:500 |
| Collagen I | Mouse | Abcam | ab6308 | 1:500 |
| Giantin (Golgi) | Rabbit | Abcam | ab80864 | 1:500 |
| Ki67 | Rabbit | Abcam | ab15580 | 1:500 |
| Cytokeratin 7 | Mouse | Abcam | ab9021 | 1:500 |
| Cytokeratin 17 | Rabbit | Abcam | ab53707 | 1:500 |
| **Secondary antibodies** | | | | |
| **Type** | | **Company** | **Cat. N°** | **Dilution** |
| Alexa Fluor® 647 Donkey anti-rabbit IgG (H+L) | | Invitrogen | A32795 | 1:500 |
| Alexa Fluor® 488 Chicken anti-rabbit IgG (H+L) | | Invitrogen | A21441 | 1:500 |
| Alexa Fluor® 568 Donkey anti-mouse IgG (H+L) | | Invitrogen | A10037 | 1:500 |
| Alexa Fluor® 405 Donkey anti-rabbit IgG (H+L) | | Invitrogen | A48258 | 1:500 |
| **Fluorescent dyes** | | | | |
| Alexa Fluor™ Plus 647 Phalloidin | | Thermo Fisher Scientific | A30107 | 1:500 |
| Alexa Fluor™ Plus 568 Phalloidin | | Thermo Fisher Scientific | A12380 | 1:500 |

**Supplementary Table 1: List of antibodies used for immunofluorescence experiments.**

**Supplementary Table 2**

| GENE NAME | PROTEIN NAME | PRIMER SENS | PRIMER SEQUENCE 5'-3' |
|---|---|---|---|
| TANGL | Transgelin | FWD | GGTGGAGTGGATCATAGTGC |
| TALGN | Transgelin | REV | ATGTCAGTCTTGATGACCCCA |
| POU5F1 | OCT4 | FWD | GTATTCAGCCAAACGACCATC |
| POU5F1 | OCT4 | REV | CTGGTTCGCTTTCTCTTTCG |
| EPCAM | Epcam | FWD | CCATGTGCTGGTGTGTGAA |
| EPCAM | Epcam | REV | TGTGTTTTAGTTCAATGATGATCCA |
| CCND1 | Cyclin D1 | FWD | TCTACACCGACAACTCCATCCG |
| CCND1 | Cyclin D1 | REV | TCTGGCATTTTGGAGAGGAAGTG |
| FN1 | Fibronectin | FWD | ATTCCAATGGTGCCTTGTGC |
| FN1 | Fibronectin | REV | TCCCACTGATCTCCAATGCG |
| ZEB2 | Zinc Finger E-Box Binding Homeobox 2 | FWD | CAAGAGGCGCAAACAAGCC |
| ZEB2 | Zinc Finger E-Box Binding Homeobox 2 | REV | GGTTGGCAATACCGTCATCC |
| SNAI1 | Snail | FWD | TCGGAAGCCTAACTACAGCGA |
| SNAI1 | Snail | REV | AGATGAGCATTGGCAGCGAG |
| TJP1 | ZO1 | FWD | GAATGATGGTTGGTATGGTGCG |
| TJP1 | ZO1 | REV | TCAGAAGTGTGTCTACTGTCCG |
| CDH2 | N-Cadherin | FWD | GTGCATGAAGGACAGCCTCT |
| CDH2 | N-Cadherin | REV | GCCACTTGCCACTTTTCCTG |
| KRT7 | Cytokeratin 7 | FWD | GGATGCTGCCTACATGAGC |
| KRT7 | Cytokeratin 7 | REV | CCAGGGAGCGACTGTTGT |
| KRT14 | Cytokeratin 14 | FWD | TGAGAAGGTGACCATGCAGA |
| KRT14 | Cytokeratin 14 | REV | ATTGTCCACTGTGGCTGTGA |
| KRT17 | Cytokeratin 17 | FWD | GGTGGGTGGTGAGATCAATGT |
| KRT17 | Cytokeratin 17 | REV | CGCGGTTCAGTTCCTCTGTC |
| KRT18 | Cytokeratin 18 | FWD | GACACCAATATCACACGACTG |
| KRT18 | Cytokeratin 18 | REV | GGCTTGTAGGCCTTTTACTTC |
| CDH1 | E-Cadherin | FWD | GTGTCATCCAACGGGAATGC |
| CDH1 | E-Cadherin | REV | TGGCGGCATTGTAGGTGTTC |

**Supplementary Table 2: Sequences of RT-qPCR primers used for gene expression analysis.**

### Supplementary legends

**Supplementary Video 1: Time-lapse imaging of MED formation.**

Bright-field imaging showing the formation of the MED's epithelium over 128 hours. Images were captured right after encapsulation, every 2 hours. Playback speed: 10 frames per second. Scale bar, 100 µm.

**Supplementary Video 2: 3D reconstruction of the MED**

3D reconstruction of the MED on day 7. The video shows a 360° rotation of the tissue segment, stained for F-actin (phalloidin, magenta) and nuclei (Hoechst, cyan).

**Supplementary Video 3: 3D reconstruction of the MESD**

3D reconstruction of the MESD on day 7. The video shows a 360° rotation of the tissue segment, stained for F-actin (phalloidin, white) 3T3-GFP fibroblasts (green) and nuclei (Hoechst, cyan).
